# Computational Investigation Of Blood Flow And Flow-mediated Transport In Arterial Thrombus Neighborhood

**DOI:** 10.1101/2020.06.11.147488

**Authors:** Chayut Teeraratkul, Zachariah Irwin, Shawn C. Shadden, Debanjan Mukherjee

## Abstract

A pathologically formed blood clot or thrombus is central to major cardiovascular diseases like heart attack and stroke. Detailed quantitative evaluation of flow and flow-mediated transport processes in the thrombus neighborhood within large artery hemodynamics is crucial for understanding disease progression and assessing treatment efficacy. This, however, remains a challenging task owing to the complexity of pulsatile viscous flow interactions with arbitrary shape and heterogeneous microstructure of realistic thrombi. Here, we address this challenge by conducting a systematic parametric simulation based study on characterizing unsteady hemodynamics and flow-mediated transport in the neighborhood of an arterial thrombus. We use a hybrid particle-continuum based finite element approach to handle arbitrary thrombus shape and microstructural variations. Results from a cohort of 50 different unsteady flow scenarios are presented, including unsteady vortical structures, pressure-gradient across the thrombus boundary, finite time Lyapunov exponents, and dynamic coherent structures that organize advective transport. We clearly illustrate the combined influence of three key parameters - thrombus shape, microstructure, and extent of wall disease - in terms of: (a) determining hemodynamic features in the thrombus neighborhood; and (b) governing the balance between advection, permeation, and diffusion to regulate transport processes in the thrombus neighborhood.

## 1 Introduction

According to latest statistics from the American Heart Association [51], cardiovascular diseases like heart attack and stroke continue to be a leading global cause of death – accounting for over 800,000 deaths annually in the United States alone. Pathological clotting of blood, also referred to as Thrombosis, is a primary cause or complication in stroke and coronary heart disease [21,55]. Thrombotic phenomena in patients are intimately related to blood flow, transport, and flow-induced forces [57,20,33,15]. Unsteady pulsatile viscous blood flow in the neighborhood of a thrombus - a pathologically formed clot - can determine the transport of platelets and coagulation proteins, as well as thrombolytic drug during therapy [14,5]. Flow-induced forces on the thrombus are known to influence thrombus volume and growth, as well as fragmentation and thrombo-embolization risks [3,12,2,17,18]. A comprehensive assessment of flow and flow-mediated transport in the thrombus neighborhood is therefore critical for understanding disease progression and thrombolytic treatment efficacy, and yet remains a challenging task. A key underlying challenge comprises understanding the interaction of unsteady pulsatile hemodynamics with realistic human thrombi of arbitrary shape and heterogeneous morphology and microstructure. Real human thrombi are known to be composite, with platelets and fibrin strands constituting primary ingredients [56,54,64,53]. The interconnected interstitial space in the thrombus interior leads to permeability of the thrombus at the macroscale, which is an important parameter for thrombus biomechanics and transport phenomena [58,4]. Several prior studies have advanced our understanding of flow and transport processes in thrombus micro-environment in microscale thrombi through experiments conducted using mouse injury models [33,54], and microfluidic flow systems [32,62]. Computer simulations have provided a viable alternative to complement experimental assays and imaging studies, having been used to model both microscale [37,59,60,45,47,61] and macroscopic [27,28,36,63] thrombus biomechanics and biotransport phenomena. Handling large artery hemodynamics interactions with arbitrary shape and heterogenous microstructures has remained a challenge in existing computational approaches. In prior work, we developed a hybrid particle-continuum fictitious domain computational method for addressing this state-of-the-art challenge [31]. This method enables easy representation of shape and microstructural features, and parametric variations thereof, by using a discrete particle description of the thrombus which is embedded weakly into a continuous background fluid mesh. The focus of [31] was on developing the computational methodology details, and demonstrating numerical features of the method; and aspects of transport phenomena mediated by the unsteady dynamic flow and pressure distributions were not explored. Here, we use the particle-continuum approach to conduct a parametric simulation study on flow and flow-mediated transport in the thrombus neighborhood within flow environments mimicking that of large artery hemodynamics. Specifically, our objectives are to: (a) characterize key features of the unsteady flow environment around arterial thrombi; (b) illustrate how key parameters like thrombus shape, microstructure, and wall leakage due to disease influence flow and transport; and (c) quantitatively establish the various ways in which flow mediates transport within and outside the thrombus.

## 2 Methods

### 2.1 Hybrid particle-continuum finite element hemodynamics model

Pulsatile viscous blood flow in the neighborhood of an arbitrary thrombus was modeled using a hybrid particle-continuum numerical approach based on a stabilized fictitious domain finite element framework we have devised in prior work [31]. Details of the formulation are not reproduced here for conciseness. Briefly, we assume an overall background fluid domain within which the thrombus is embedded (see Figure 1), instead of meshing the fluid and the thrombus as separate domains with a mesh that conforms with the thrombus boundary. Assuming that blood is a Newtonian fluid, we solved the Navier-Stokes equations for fluid momentum balance, and continuity equation for mass balance, using a Petrov-Galerkin stabilized finite element formulation [7,16]. The variational form for this finite element formulation is given as follows:

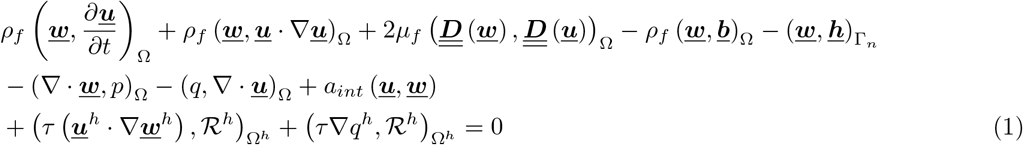

where (*a, b*)Ω ≡ ∫ *a · b d*Ω; Ω is the overall computational domain; Ω^*h*^ represents the discretized computational domain; *ρ_f_* is blood density; *μ_f_* is blood viscosity; <u>***u***</u> and *p* are the flow velocity and pressure; <u>***w***</u> and *q* are the respective velocity and pressure test functions; 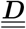 denotes the strain rate tensor; and 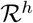 is the residual of the momentum and mass balance equations. The last two terms in Equation 1 represent contributions from Petrov-Galerkin stabilization for convective phenomena and linear pressure-velocity interpolation respectively. The stabilization parameter *τ* was chosen as specified in prior work [31]. Contributions from resistance and Windkessel based boundary conditions, typical in vascular hemodynamics modeling, were accounted for through <u>***h***</u>. For example, a resistance boundary condition (with resistance value *R_m_*) over a part of the boundary Γ_*m*_ can be formulated as follows:

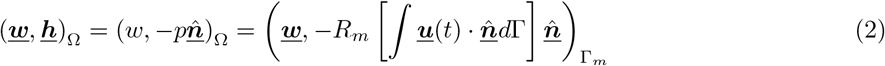

and we refer to [50] and [23] for further relevant details regarding mathematical formulation of such boundary conditions. For the fictitious domain formulation [31], coupled interaction between the thrombus domain and the flow domain was handled using custom coupling terms a_*int*_ (<u>***u**, **w***</u>) in Equation 1. Here we employed the penalty-function approach formulated in [31], where the fluid velocity (<u>***u***</u>) is constrained to take-up the local velocity (<u>*v*</u>_0_) within the thrombus domain (Ω_T_):

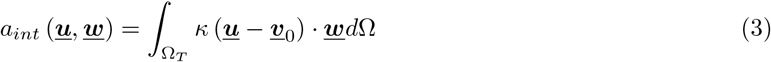

with *k* being a penalty parameter chosen based on element size *h* locally as follows:

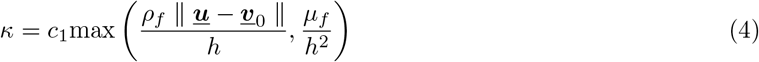

**Figure 1:**
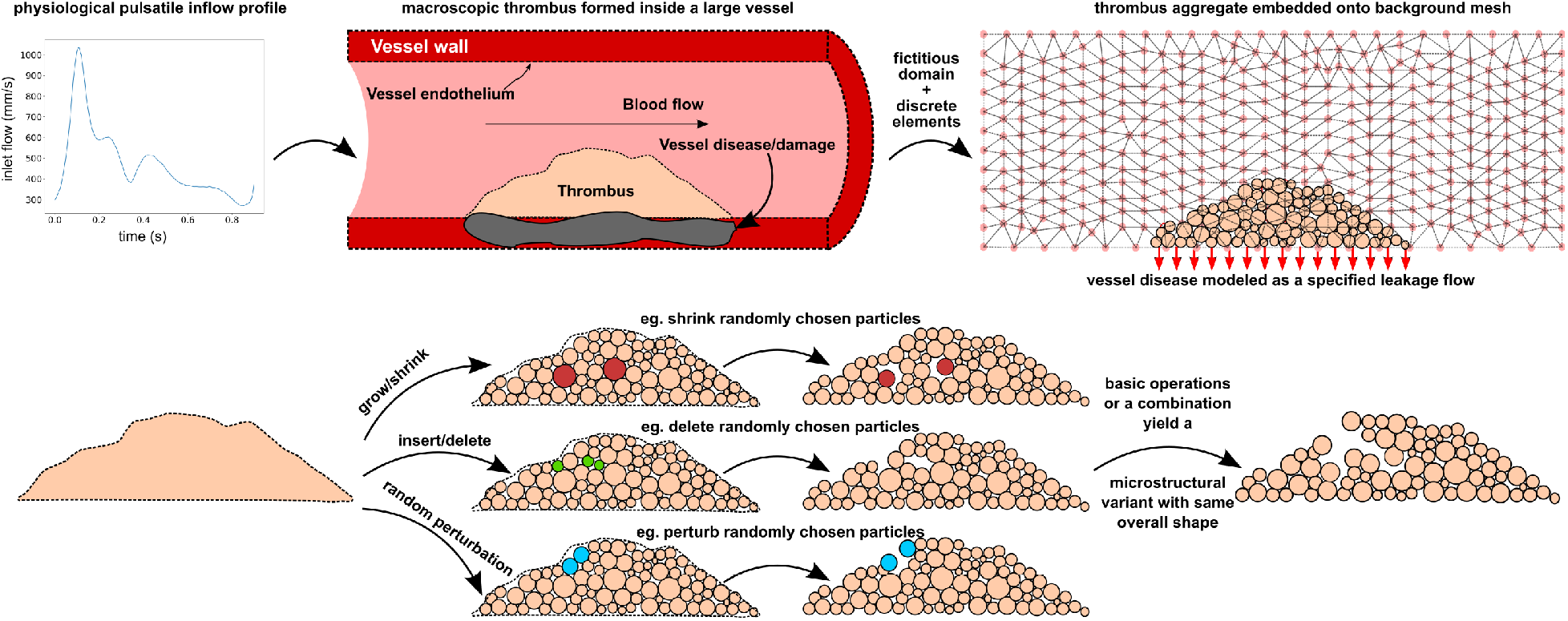
A schematic illustration of the various conceptual steps involved in the hybrid particle-continuum fictitious domain framework. The top panel illustrates the pulsatile inflow data, simulation domain configuration, and the fictitious domain approach with leakage boundary conditions to model wall disease. The bottom panel provides a cartoon illustration of the discrete particle based microstructural variation modeling.

In this study, we used *c*_1_ = 500.0 for all simulations. In accordance with the hybrid particle-continuum approach, the thrombus domain Ω_T_ was modeled using a collection of mesh-free, off-lattice, discrete elements 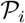 such that: 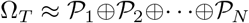. Within this discrete element domain, <u>***v***</u>_0_ then denotes the local velocity of the discrete element itself, which would be 0 for a stationary discrete element domain. This approach has been demonstrated to be effective at handling the arbitrary shape and heterogeneous microstructure of physiologically realistic thrombi [31]. While the original formulation presented in [31] models each discrete element 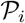 as a generalized superquadric geometry, for the purpose of this study, we modeled 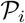 as individual spherical particles. For each element, we used the sphere definition, to identify whether a finite element quadrature point is *‘inside’* or *‘outside’* the thrombus domain. For all points located *‘inside’* - the fictitious domain penalty contributions were assembled into the global matrix system for the fluid flow problem.

### 2.2 Modeling thrombus microstructure variations

We used the framework devised in prior work [31] to reconstruct a thrombus using discrete elements. Briefly, this framework relies on post-processing of thrombus image data (medical images or microscopy images); identification of thrombus manifold geometry via image-segmentation; identification of additional spatial composition information if available; and feeding the thrombus data into a geometric tessellation based algorithm to create a discrete particle reconstruction of the thrombus domain. The resultant reconstruction represents a coarsened, mesoscopic approximation of the thrombus internal domain, with an effective mesoscale porosity and pore-space network. The discrete element reconstruction enables flexible modeling of a range of microstructural variations in the thrombus interior. One approach is to modify the shape of each discrete element in a parametric manner, as described in [31]. Here, we devised a set of additional basic operations to account for a wider range of microstructures (see for reference Figure 1). These operations involved: (a) growing or shrinking a randomly selected subset of discrete elements by a uniform random growth/shrinkage factor; (b) inserting or deleting discrete elements from a randomized selection of locations; (c) perturbing the location of a randomly selected subset of discrete elements. A set of arbitrary sequences of growth-shrinkage, insertion-deletion, and perturbation operations were used on a baseline discrete element reconstruction from microscopy image data, to generate a range of microstructural variants for the same thrombus. These microstructural variants of the discrete element reconstruction were constrained to keep the overall shape and size of the thrombus fixed, and the overall statistical distribution of the particle sizes across the variants remained the same.

### 2.3 Modeling of diseased wall state

Diseased vessel wall status was modeled conceptually in form of a pervious or leaky wall at the base of the thrombus. This rationale was based on discussions in existing literature such as [48]. Extent of leakage was scaled with respect to the total incoming volumetric flow at the channel inlet as shown below:

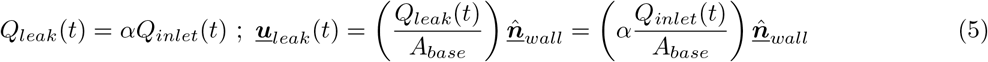

where *Q_inlet_* is the total inlet flow rate; *A_base_* is the area of the wall along the base of the thrombus (that is, diseased wall area); and 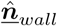 is the outward normal from the vessel wall at the thrombus site. The average velocity derived from this leakage flow rate was then imposed along the vessel wall at the base of the thrombus as a time-varying Dirichlet boundary condition within the fictitious domain finite element formulation. The concept is illustrated in Figure 1. This simple method has few distinct advantages. First, the extent of wall disease or damage is characterized by a single parameter *α* which has a direct phenomenological interpretation. Second, the methodology is easily applicable in both 2D and 3D geometries, enabling extension to real arterial geometries as long as the clot location is identifiable (for example, using image data). Furthermore, the dynamic leakage flow in real thrombus sites is dependent upon pressure and permeability of the extravascular space [48]. Detailed data on these extravascular parameters are needed for refined estimates of leakage flow extent and dynamics. If this data is available, our approach here can be extended to incorporate the data in form of spatiotemporally varying leakage flow estimate modeled as a single spatiotemporally varying model parameter *α* (*<u>**x**</u>,t*); which provides added numerical convenience in reducing the number of model parameters.

### 2.4 Lagrangian computation of Finite Time Lyapunov Exponent (FTLE) field

For a given flow velocity field data, we adopted a Cartesian grid based tracer integration approach to compute the Finite Time Lyapunov Exponent (FTLE) field as devised extensively in prior works by Shadden et.al. [43,42,40]. Specifically, a Cartesian grid of massless Lagrangian tracers were seeded in the interior of the flow domain. For our fictitious domain approach, the Cartesian grid included the interior of the fictitious thrombus domain. The position of each tracer was integrated based on the flow velocity interpolated at the location of the tracer at each instant as follows:

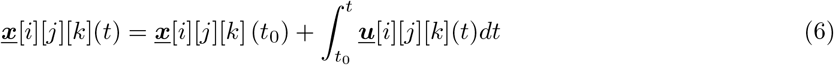

where we have used the notation [*i*][*j*][*k*] to denote indexing based on the grid used for the computation. The integration in Equation 6 was evaluated numerically using a one-step explicit four stage Runge-Kutta method. As discussed in several published works, locating the cell/element where the tracer resides at a given instant for integration of its velocity can be a non-trivial and time-consuming task, especially for unstructured meshes of complex geometries. Here, we mitigate this complexity by using a cell-walking algorithm for simplicial elements (triangles and tetrahedra) as proposed in [22], which relies on an algebraic check based on the tracer coordinates and element node coordinates. Considering the original seed coordinates *<u>x</u>*[*i*][*j*][*k*] (*t*_0_) as the reference configuration, and the mapped coordinates due to the flow at time *t* - <u>*x*</u>[*i*][*j*][*k*] (*t*) - as the current configuration, we defined a deformation gradient 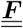 for the tracer kinematics as follows, based on the formulation presented in [40].

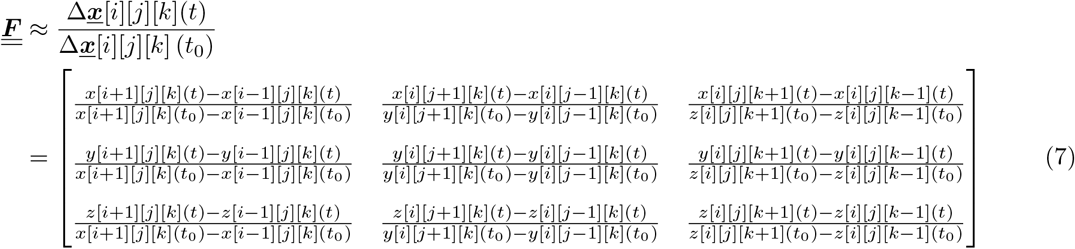

The right Cauchy-Green deformation tensor was then computed as follows:

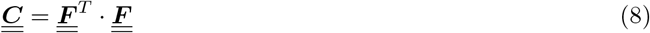

and subsequently, the FTLE field values were calculated as a scaled version of the natural logarithm of the square root of the maximal eigenvalue of the right Cauchy-Green deformation tensor as follows:

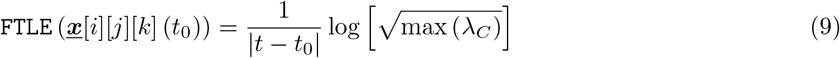

with λ_*C*_ denoting eigenvalues of the tensor 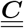. The computed FTLE field was mapped back onto the original Cartesian grid where tracers were seeded (hence the notation on the left hand side of Equation 9).

### 2.5 Extraction of FTLE ridges

Ridges of the FTLE field define coherent structures that organize advective mass transport [43]. For this study, we computed the FTLE field over a 2-dimensional Cartesian structured grid, and interpreted them as pixels of an image for identifying ridges in the transformed FTLE pixel intensity data. Using the Scikit-Image [49] library in Python, we computed 2-dimensional Hessian matrix of the resulting FTLE-field image pixel intensity distribution, and the corresponding eigenvalues of the pixel intensity Hessian matrix. This was followed by a thresholding of pixels based on the computed eigenvalues using the Otsu method [35,29] to identify the ridges in the images, quantified in whole pixel units. FTLE fields were pre-processed into portable natural graphics (png) formatted images at high dpi values, and cropped to field view, prior to employing these image-based operations.

### 2.6 Model system and design of experiments

The model system for this study comprised a 2-dimensional rectangular channel domain with width equivalent to that of the human common carotid artery (≈ 6.0 *mm*), and length equal to 5 times the width. Blood was modeled as a Newtonian fluid with constant density *ρ_f_* = 1.06 *g/cc*, and bulk viscosity *μ_f_* = 4.0 *cP*. A pulsatile inflow profile was specified at the channel inlet, based on measured common carotid artery flow profile data available in literature [26]. A set of realistic thrombus morphology images obtained from human whole blood clotting experiments as reported in [12] was processed using the framework described in Section 2.2. The thrombus aggregate dimensions were scaled up to arterial scales while preserving their overall shape and aspect ratio. Using the techniques outlined in Section 2.2 a set of 6 microstructure models were generated for each of the 2 clot specimen in [12], leading to 12 thrombus models as illustrated in Figure 2. With statistically similar particle size distribution, the difference between maximum and minimum porosity across all microstructures for both models was around 12.5 percent. For each microstructure model, four different scenarios for diseased wall status were considered corresponding to *ρ* = 0.0,0.1,0.2,0.4 - that is, 0%, 10%, 20%, and 40% leakage flow across the diseased wall respectively. We remark here that the case with *ρ* = 0.0 will also correspond to microfluidic devices with fixed impermeable channel geometries. Despite being a relatively high leakage fraction, *ρ* = 0.4 was included to quantitatively demonstrate how the flow would behave for a significant amount of leakage. In addition, for each of the two clots, a separate case was considered where the clot boundary was modeled as a purely rigid, impermeable, no-slip boundary wall. For each of the 50 resulting cases, unsteady hemodynamics was simulated for three cardiac cycles using the fictitious domain finite element method (Section 2.1). A two-step meshing strategy was employed with a finer mesh sizing imposed on a refinement region around the thrombus, and a coarser sizing imposed on rest of the flow domain away from the thrombus. Target element size for coarser region was around 150 microns, guided by mesh refinement analysis (see [31]), and elements in the refinement region were 3 times smaller. Subsequently, FTLE fields were computed using the flow velocity data from the final cardiac cycle using the tracer integration approach in Section 2.3.

**Figure 2:**
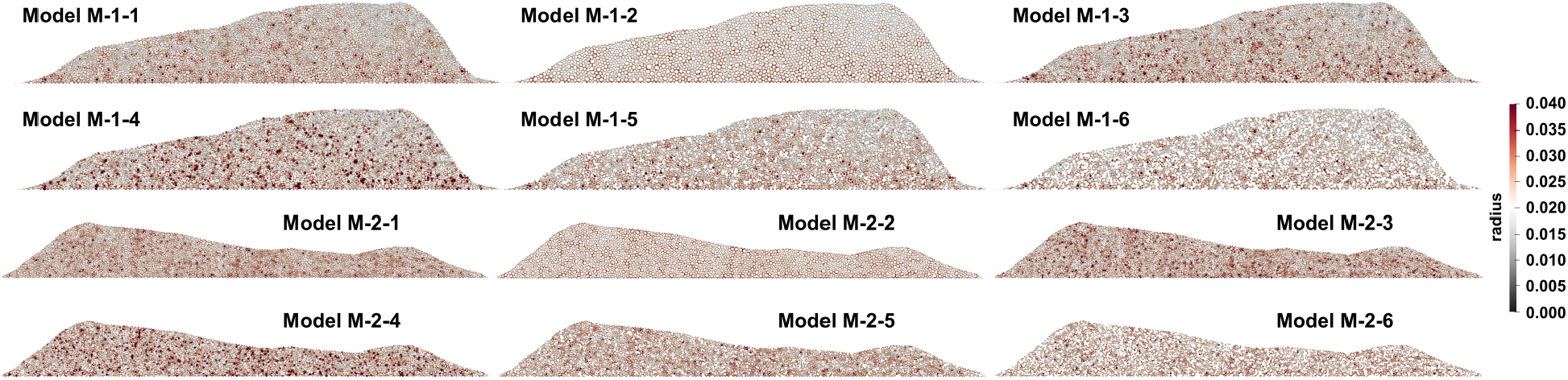
Illustration of all the 12 different thrombus models, derived from a combination of 2 thrombus model shapes (based on data in [12]) and 6 microstructural variations per model. Models are indexed using a naming scheme: M-i-j; where i indicates the shape, and j indicates the microstructural variant. All models are colored by the size of the discrete elements or particles used to represent the microstructures.

## 3 Results

### 3.1 Unsteady hemodynamics in the thrombus neighborhood

Computed unsteady flow patterns were visualized for the final cardiac cycle for each of the numerical experiments undertaken for this study. Flow patterns were visualized using surface line integration convolution (LIC), which is a texture advection based visualization technique for vector fields that clearly demarcates local rotational or vortical regions (vortex cores) [8] in addition to velocity magnitudes (based on LIC color). Figure 3 presents LIC patterns colored by velocity magnitude for varying extents of wall leakage (*α*) at peak systole for thrombus models M-1-X. Figure 4 presents the same for models M-2-X. Additional figures presenting flow LIC visualizations for mid-systolic acceleration, peak systole, and mid-systolic deceleration (*refer Figure 1 for flow profile*) are provided for both sets of thrombus models in the Supplementary Material (see *Figures S1 and S2 as well as supplementary animations*). We observe that, for fixed clot shape and fixed wall leakage parameter, the larger scale hemodynamic patterns, vortical structures, and recirculation features show minimal variations across the microstructural variants considered. However, variation in the extent of wall leakage significantly affects the unsteady hemodynamic patterns in the thrombus neighborhood. To further illustrate, and support, this observation, we quantified the difference in velocity fields by comparing the flow velocities around M-1-X and M-2-X to those obtained from unsteady flow computations around a purely rigid and impermeable thrombus boundary (modeled as a hole in the domain with boundaries matching the thrombus manifold geometry). The differences in local flow velocity magnitudes were integrated across the entire fluid domain outside the thrombus, and the time-varying integrated velocity differences for models M-1-X are presented in Figure 5 (*equivalent plot for M-2-X models included in Supplementary Material Figure S3*). As seen in the time varying differences, as well as the maximum values for each microstructure-leakage combination (inset), variations across the different microstructures are significantly small (*see also Supplementary Material Figure S4 for a visualization of the pressure dynamics at the leakage site*.) Together, these results clearly indicate how thrombus shape, microstructure, and wall condition can influence macroscale unsteady flow. We observe that: (a) microstructure leads to minor variations in large scale flow patterns; and (b) to a leading order, overall thrombus shape and wall conditions have a significantly greater influence on larger-scale flow structures than finer thrombus microstructure. These trends will understandably change for cases where extreme microstructural variants are considered (eg. a very dense structure compared against a very sparse structure), which was not the focus of these experiments.

**Figure 3:**
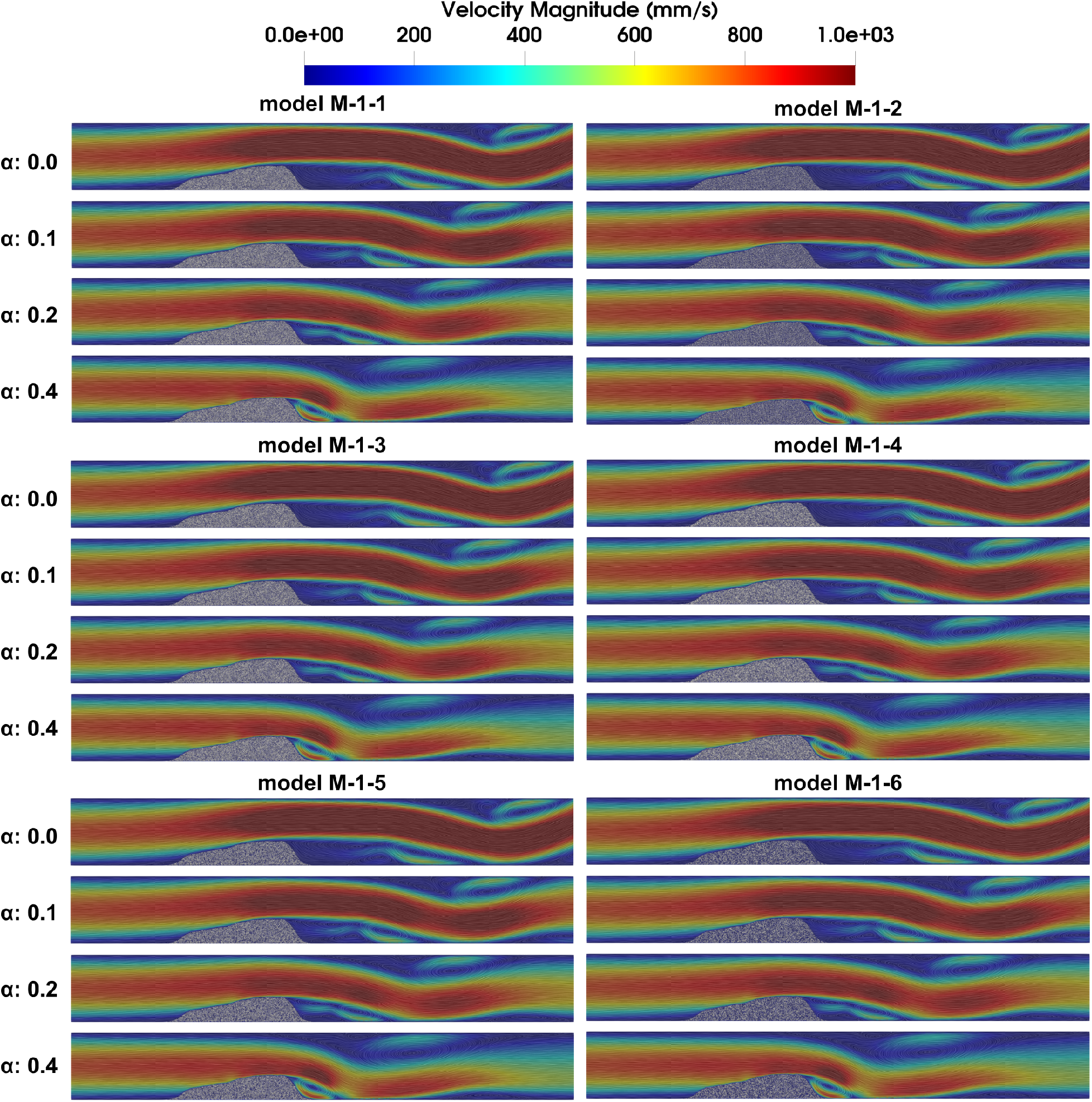
Flow velocity fields at peak systole for all parametric combinations for thrombus model 1 (that is, model id’s M-1-x, x=1 – 6), visualized using line integration convolution (LIC) maps colored by velocity magnitude. For each thrombus shape-microstructure combination (M-1-x), the wall leakage parameters *α* increases from top to bottom.

**Figure 4:**
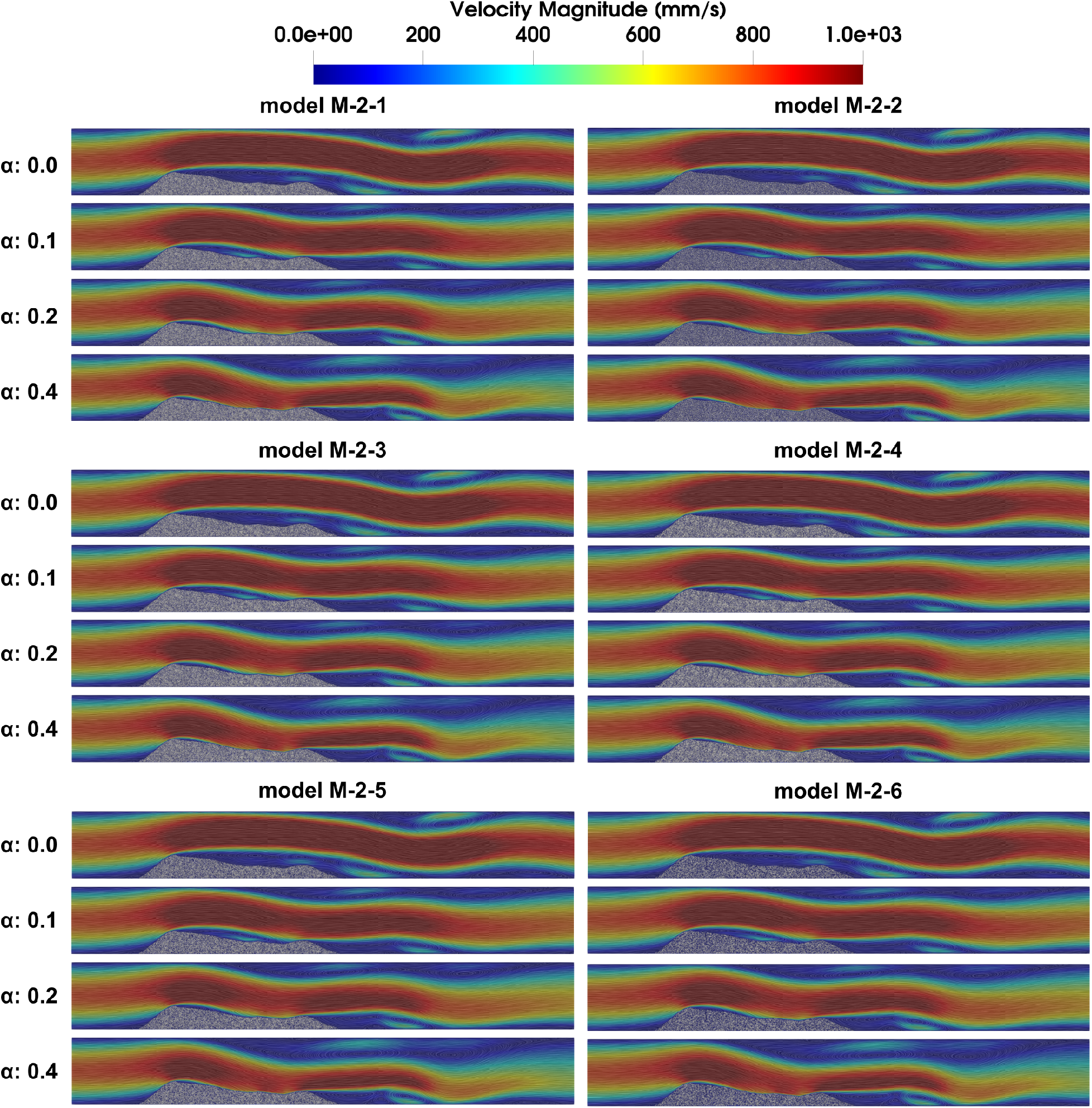
Flow velocity fields at peak systole for all parametric combinations for thrombus model 2 (that is, model id’s M-2-x, x=1 – 6), visualized using line integration convolution (LIC) maps colored by velocity magnitude. For each thrombus shape-microstructure combination (M-2-x), the wall leakage parameters *α* increases from top to bottom.

**Figure 5:**
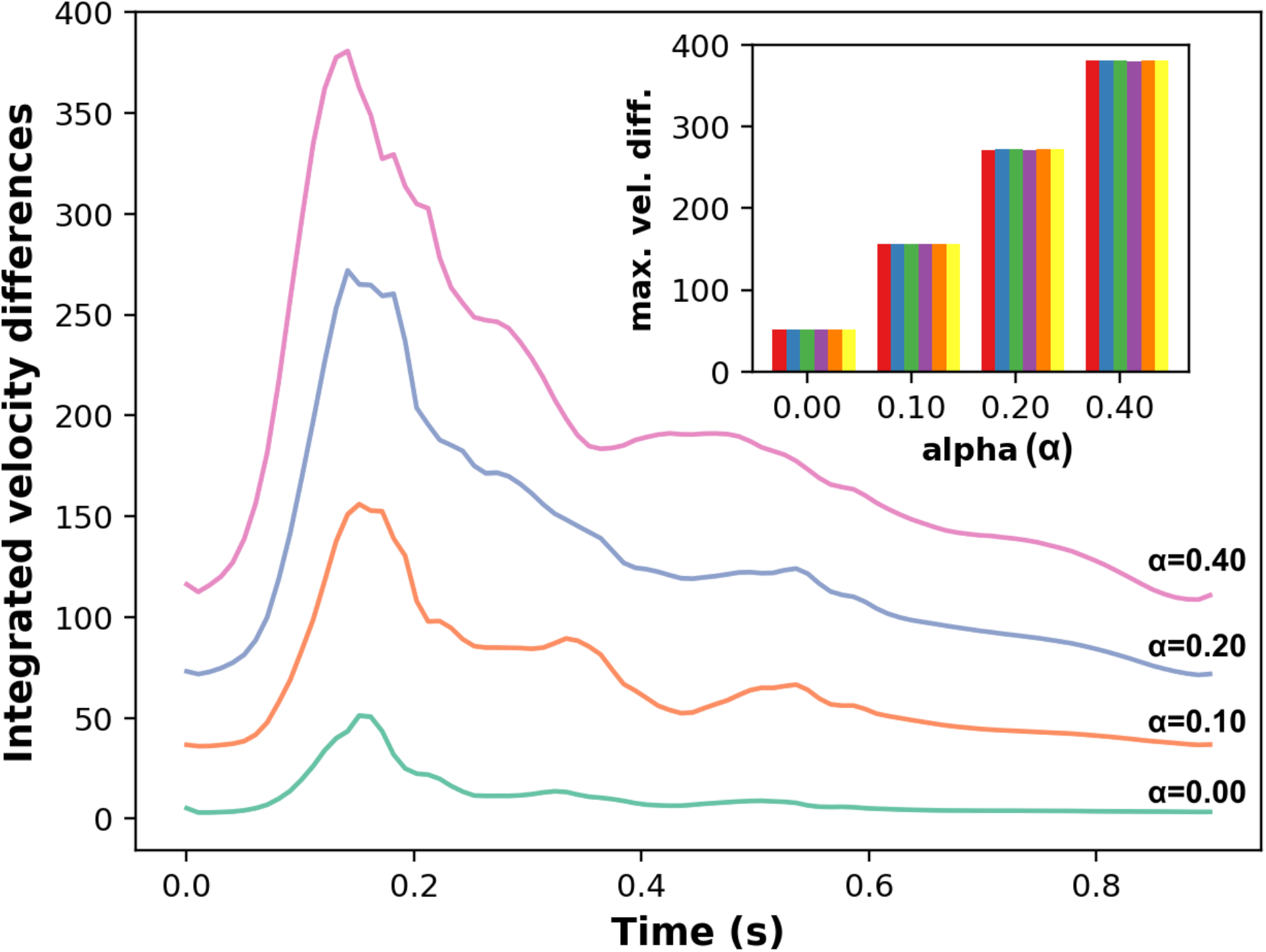
Differences in flow velocity fields around models M-1-x computed with respect to flow velocities around a rigid thrombus with impermeable walls and integrated over the entire flow domain. The main visual illustrates the variation in this integrated difference over time in one cardiac cycle for varying leakage parameter *α*. The curves represent mean values computed across the 6 microstructural variants. Inset visual illustrates the maximum differences for varying leakage parameters across all microstructural variants, confirming the minor influence that microstructure has in comparison with wall leakage in affecting flow around a given thrombus shape. All data are in units of velocity mm/sec.

### 3.2 Pressure gradient across the thrombus boundary

Pressure gradient across the thrombus domain is an important driver for flow-mediated permeation into the thrombus interior. Based on velocity and pressure data for each of the numerical experiments, we computed pressure gradient across the thrombus boundary, in a direction normal to the boundary, at successively varying locations along the thrombus geometry. We refer to this here as the thrombus boundary pressure gradient or TBPG. Figure 6 illustrates the normalized values of computed TBPG - averaged in time over one cardiac cycle - plotted along the longitudinal span of the thrombus domain, for varying microstructures and varying wall leakage parameter (*α*). Positive values of TBPG here indicate a pressure gradient favoring flow permeation and flow-mediated transport into the clot. Negative values, on the contrary, denote adverse pressure gradients for flow permeation. The time-averaged TBPG was integrated along the fluid-thrombus interface boundary for every combination of shape, microstructure, and wall leakage parameter. Resulting data for thrombus models M-1-X and M-2-X are presented in Figure 7. Increase in extent of wall leakage, in the absence of any other thrombus structure modifications, is seen to increase the pressure gradient across the thrombus. For each wall leakage parameter, we computed the mean and coefficient of variation of the resultant integrated time-averaged TBPG value across all microstructures. This is further illustrated in Figure 7. Together, Figures 6 and 7 show the combined influence of thrombus shape, thrombus microstructure, and wall leakage on pressure-gradient across the thrombus, and the resultant potential for flow permeation in the unsteady arterial hemodynamic environment. The TBPG, and resulting pressure-driven permeation, is non-uniform along the thrombus boundary, which is in agreement with prior studies [13,5]. We observe that thrombus shape influences the locations and extent of positive TBPG significantly. Specifically, at points along the geometry where the viscous flow separates (*marked in Figure 6 with red dotted lines*) we observe a sharp change in pressure gradient. This agrees with expectations based on fluid mechanical considerations of viscous boundary layer and flow separation. Additionally, the effect of microstructural variations are more visible in the resultant TBPG values than in the resultant large-scale flow patterns. Specifically, from Figure 7, we observe that the coefficient of variation in TBPG values across all microstructures is an order of magnitude higher for cases where *α* = 0 (that is, no wall leakage) when compared to cases with non-zero *α*. For all cases with non-zero *α*, the variability in TBPG increases with increasing *α*. These observations indicate that the thrombus microstructure has a relatively more noticeable effect on the extent of flow-mediated permeation into the thrombus as compared to the large scale flow structures around the thrombus.

**Figure 6:**
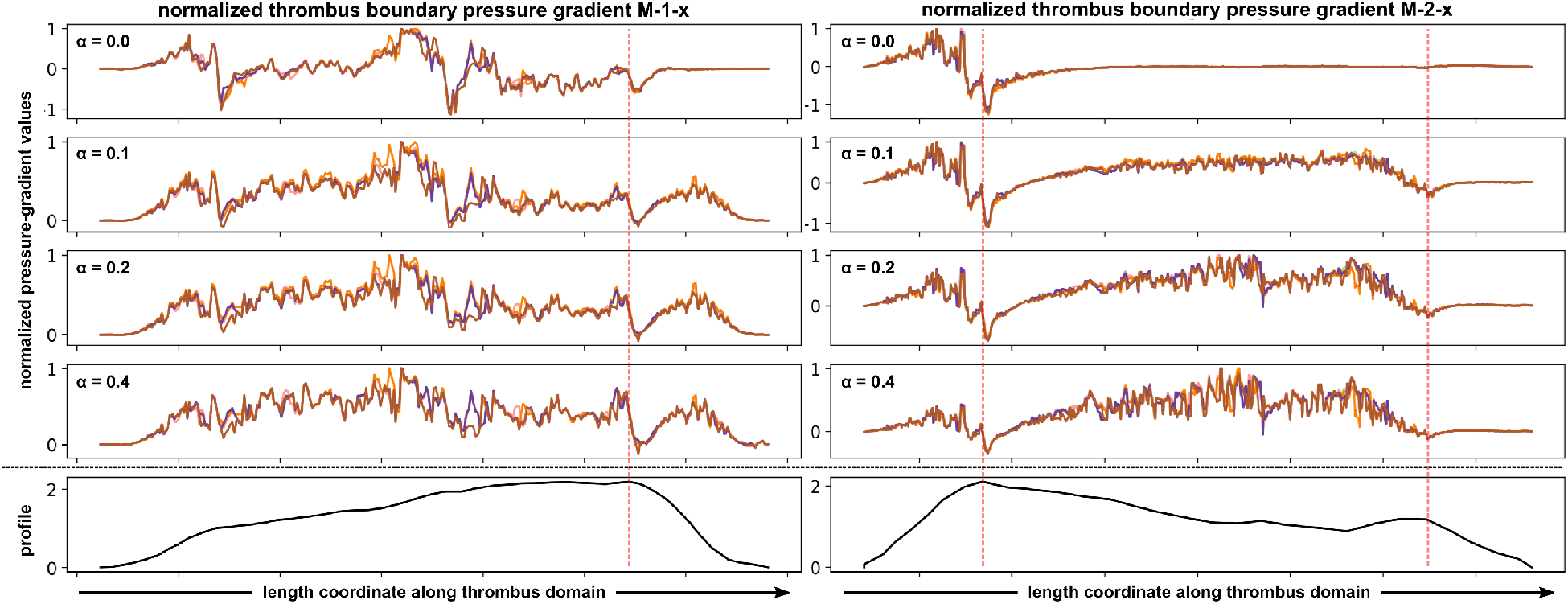
Normalized values of pressure-gradients normal to the thrombus boundary, computed along the entire boundary length for varying wall leakage parameter *α*. Each plot shows the variation of the pressure gradient with varying microstructures. The two bottom columns represent the overall thrombus shape. Red dotted lines indicate locations of expected flow separation. Positive normalized pressure gradient values favor permeation of flow into the thrombus domain.

**Figure 7:**
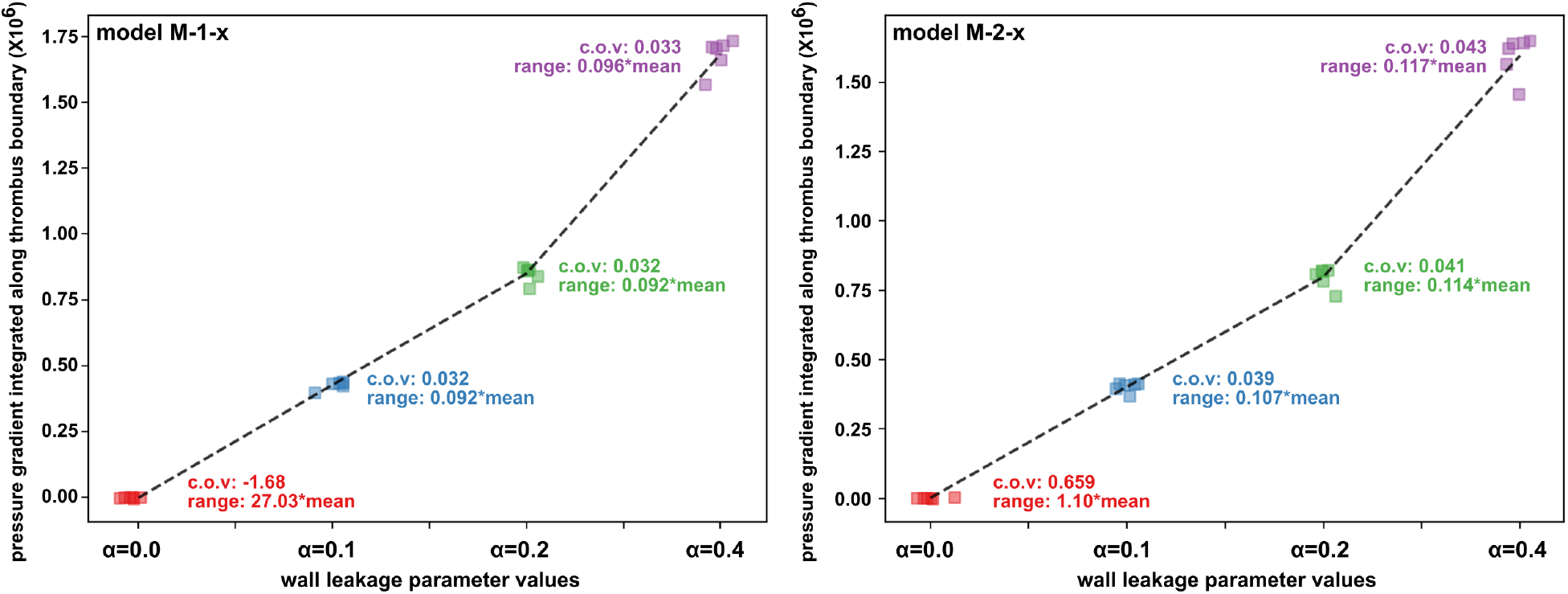
Variation of thrombus boundary pressure gradient integrated along the boundary, visualized against varying microstructures, and varying wall leakage parameter *α*. Data for thrombus models M-1-x are shown on the left, and those for models M-2-x are shown on the right.

### 3.3 Dynamic FTLE fields and coherent structures

The Finite Time Lyapunov Exponent (FTLE) values and the corresponding ridges in the FTLE scalar-field were computed as outlined in Sections 2.4 and 2.5 respectively, using velocity fields from each simulation case. In accordance with flow velocity around the thrombus, the FTLE data showed minimal variations across varying microstructures when the thrombus shape and wall leakage parameter (*α*) were held fixed. Hence, resulting FTLE field and FTLE ridges, are presented for varying clot shape and leakage for one of the thrombus microstructure models (that is, M-1-1 and M-2-1) in Figures 8 and 9 respectively. The figures illustrate the FTLE data at three instants in the cardiac cycle - mid-systolic acceleration, peak systole, mid-systolic deceleration - thereby illustrating the inherently dynamic evolution of the coherent structures forming around the thrombus. The FTLE data presented in these figures are not scaled by integration time for easy identification of the ridges. In all cases, the thrombus boundary itself constitutes a ridge. The presence of some non-zero FTLE data in the thrombus interior is, for these cases, an artifact induced due to the fictitious domain formulation and non-zero velocities in the interior domain, and does not change the interpretation of the resulting FTLE data external to the thrombus. We observe pronounced ridges forming both proximal and distal to the thrombus domain. The ridge structures formed proximal to the thrombus widens with increasing wall leakage (*α*). Distal to the thrombus, ridge structures formed follow the flow separation and reattachment dynamics, and in general get compressed closer to the thrombus domain with increasing wall leakage. Overall, a continuous enveloping FTLE ridge structure is seen to surround the thrombus in all cases, running along the thrombus length from the proximal to the distal end. Resulting flow separation and vortex dynamics also leads to pronounced FTLE ridges forming on the channel wall away from the thrombus at the distal end. The unsteady structures formed by the FTLE ridges demonstrate similarity to unsteady FTLE patterns around arterial stenosis model presented in [41], which is expected based on considerations of viscous flow separation at narrowed segments of the artery as also observed in current study. We also note that these structures show marked differences in extent and complexity from those obtained around microscale thrombi as shown in [60] (one of the first studies to compute FTLE and coherent structures around a thrombus). FTLE ridges constitute pockets and barriers for advective transport phenomena [43,19]. These results therefore indicate that the unsteady pulsatile flow, upon encountering a realistic thrombus, will generate dynamic coherent structures that organize mass transport in the thrombus neighborhood. This mass transport organization depends upon thrombus shape and size, as well as state of wall disease. In addition, it has been shown in prior work [41] that coherent structures formed by FTLE ridges are locations where integrated strain on blood-borne elements (eg. platelets) is maximized. This has key implications in mechanical platelet activation phenomena, which subsequently influences further thrombus growth and thrombosis disease progression.

**Figure 8:**
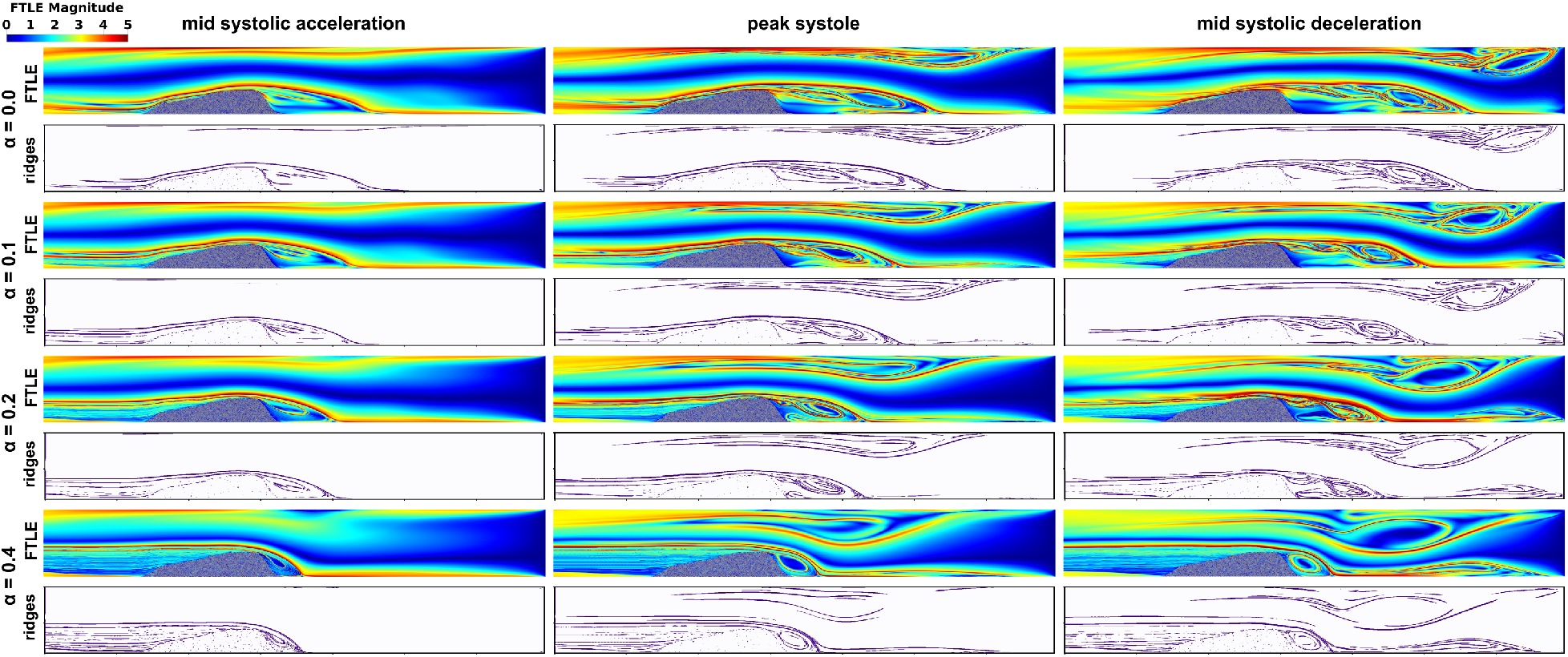
Unsteady finite time Lyapunov exponent (FTLE) fields and coherent structures formed by FTLE ridges visualized for thrombus model 1 for increasing wall leakage parameter values. To illustrate the dynamic field, data at three instances in the cardiac cycle - mid systolic acceleration, peak systole, and mid systolic deceleration are presented. All data are from simulations with case M-1-1.

**Figure 9:**
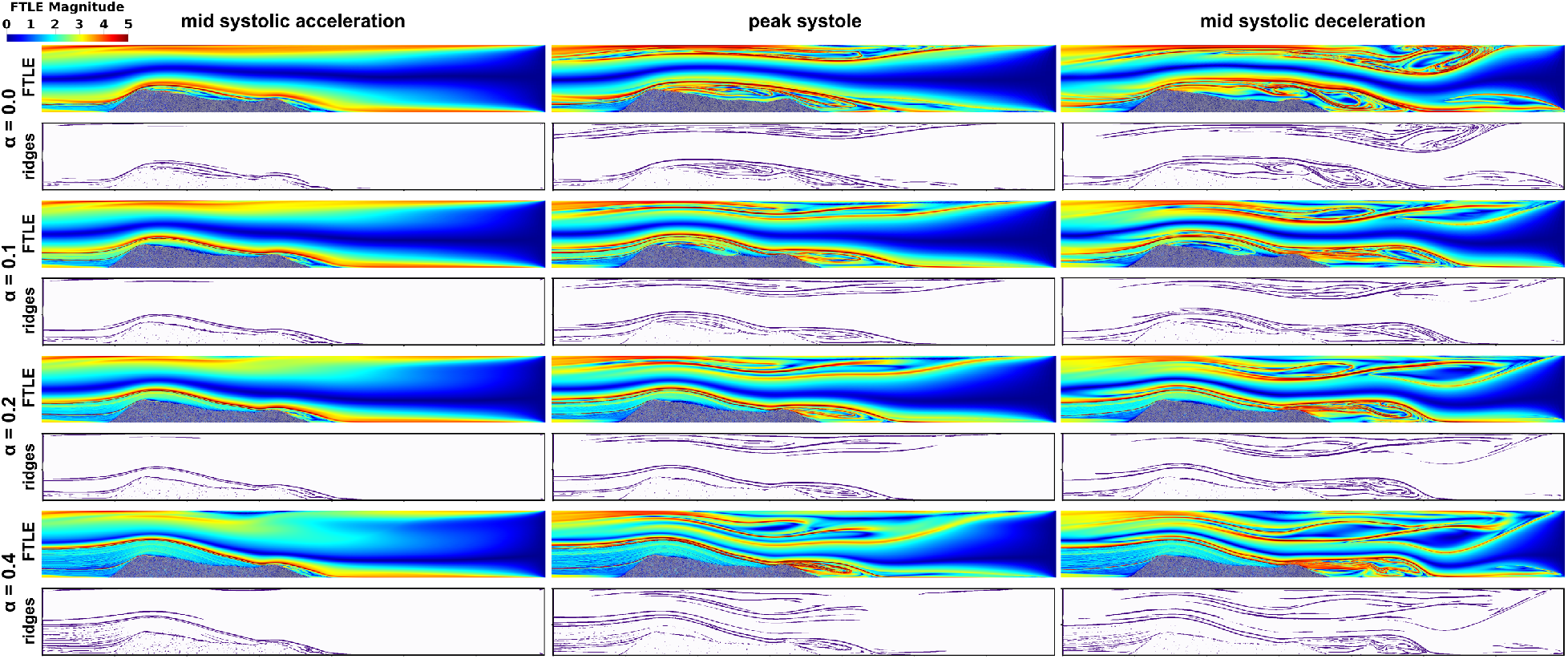
Unsteady finite time Lyapunov exponent (FTLE) fields and coherent structures formed by FTLE ridges visualized for thrombus model 2 for increasing wall leakage parameter values. To illustrate the dynamic field, data at three instances in the cardiac cycle - mid systolic acceleration, peak systole, and mid systolic deceleration are presented. All data are from simulations with case M-2-1.

## 4 Discussion

### 4.1 Coupled interplay of factors characterizing near-thrombus environment

In this study, we have characterized the thrombus microenvironment using three parameters: shape, microstructure, and wall disease. Our simulations illustrate the coupled interaction of these parameters in determining unsteady flow and flow-mediated transport in the thrombus neighborhood. Specifically, wall disease state can influence the extent of flow leakage from the thrombus site, which in turn influences the pressure gradient across the thrombus. This further influences the unsteady flow around thrombus boundary and the resulting larger scale vortex generation and recirculation. As we illustrated here, these unsteady flow features lead to formation of local pockets and barriers for transport. Variations in vortical structures with varying wall leakage can be interpreted using known fluid mechanical theories of viscous boundary layers evolving over porous surfaces with suction [6,46,39]. Presence of wall leakage draws flow in to the clot across the permeable wall, which mimics flow suction from the viscous boundary layer. Removal of flow from the boundary layer thins and stabilizes the layer, resulting in delayed or reduced extent of flow separation [39]. This is in accordance with the observations from our numerical experiments. We remark here that our study focuses on phenomena in the neighborhood of a thrombus that has already formed, and is stable. Arterial thrombi can be unstable and fragment or embolize due to flow-induced forces [18], a phenomena that we are currently investigating. During thrombus formation and growth, the synergistic interplay of the flow and wall-disease state also constitutes a crucial factor. Wall disease state influences the trans-thrombus pressure gradient, which affects flow and shear, thereby influencing the microstructure and shape of the thrombus [5,48] as it grows.

### 4.2 Assembling a flow physics concept map for thrombus neighborhood

This simulation study provides detailed insights into the various flow physics aspects that govern processes in the thrombus neighborhood in arterial flow environments. The various results and interpretations can be synthesized into a unifying concept map that elucidates flow-mediated transport phenomena in the thrombus neighborhood. We have illustrated this concept map in Figure 10. The Peclet number is a key non-dimensional descriptor for mass transport defined as *P_e_* = *UL/D* for a flow with characteristic velocity *U*, length-scale *L*, and a species with mass diffusion coefficient D. The Peclet number describes the ratio between extent of advective and diffusive mass transport. As shown here, pulsatile, viscous flow interacting with realistic thrombus shapes lead to fast flow with complex vortical structures around the thrombus. These flow structures in turn result in coherent manifolds or barriers that organize advective mass transport outside the thrombus. These higher velocity flow structures result in high Peclet number advection-dominant transport phenomena outside the thrombus. Pressure-gradient induced by the flow at the thrombus boundary drives permeation across the thrombus into the thrombus interstices. As demonstrated in our prior work [31], and other studies [24,54,30], once permeation brings flow and biochemical species into the thrombus, the small interstitial spaces in the microstructure substantially slows the flow, generating a low Peclet number diffusion dominant transport regime. Flow-mediated transport in thrombus neighborhoods thus comprise a combined interaction of advection, diffusion, and permeation. This interplay is strongly influenced by the boundary and the trans-boundary pressure gradient. Wall disease state may induce leakage, thereby changing the boundary pressure gradient, and in turn influencing the advection-diffusion-permeation interplay that governs flow-mediated transport. We remark that this concept map illustrates the general trends for how flow mediates transport of key biochemical species in the thrombus environment. Exact spatiotemporally varying concentration fields of specific constituents such as coagulation proteins can be computed using the advection-diffusion equations based on the obtained flow velocity fields, which was not the focus of this study.

**Figure 10:**
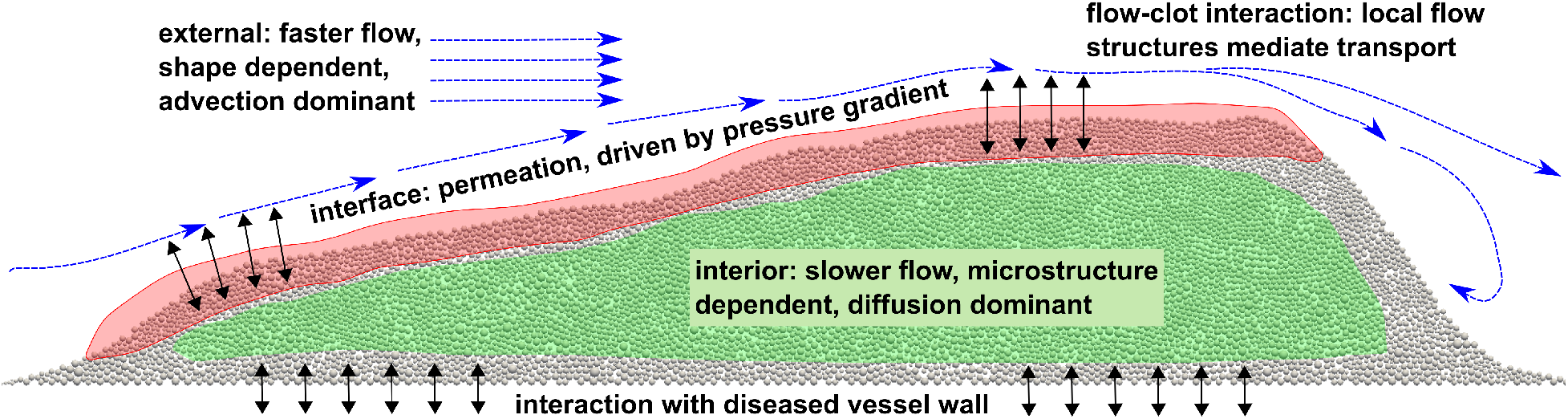
A schematic concept map derived based on data and observations form this study, that unifies the role of microstructure, shape, and wall disease or leakage in determining the three modalities of flow-mediated transport with and around a thrombus in an arterial hemodynamic environment - advection, diffusion, and permeation.

### 4.3 Performance of the fictitious domain framework

While mathematical details of the fictitious domain methodology have been outlined in prior work [31], we briefly discuss three key performance aspects here. First, the fictitious domain contribution is abstracted as a sequence of mutually independent inside-outside checks based on discrete particle data. This operation can be directly parallelized based on broadcast communication of discrete particle data across multiple processors over which the mesh is partitioned. Additionally, the background mesh is significantly less complicated than the case where each individual thrombus pore space is explicitly resolved. This makes mesh partitioning and load balancing substantially less complicated, and native partitioning and parallelization capabilities of solver libraries can be easily availed for implementing our proposed method. This simulation study extensively leveraged this ease of parallel implementation using the open source finite element library FEniCS [1]. Second, the performance of the method is intimately connected to the mesh size relative to discrete element size. Theoretically, the mesh size *h_m_* should entirely resolve the discrete element size *h_d_* (diameter of the particles). For applications with heterogeneous embedded structures (as described here), having significant variability in both *h_m_* and *h_d_*, this may lead to excessively small elements and high computational cost. Hence, we formulated a statistical approach to resolving the discrete elements, where instead of the exact values, the probability densities (pdf’s) of the two sizes are considered, and a meshing strategy is employed that: (a) has low overlap between the pdf’s for *h_m_* and *h_d_*; and (b) ensures that the mode of the pdf for *h_m_* is less than that for *h_d_*. The costs with excessively fine mesh can be mitigated also by interpolating the discrete element inside-outside functions onto a background grid (see *Figure S6 for an illustration of the interpolated discrete element inside-outside functions*), and using this interpolated function to estimate matrix contributions. Finally, and related to the previous point, the mathematical details of the fictitious domain interaction term *a_int_*(<u>***u, W***</u>)*ω* in Equation 1 needs some attention. This interaction term *a_int_* resides only on the space of the quadrature points during the finite element assembly operation; and during integration, *a_int_* is evaluated at each quadrature point directly. However, the integration can also be performed by evaluating *a_int_* at element nodes and using Galerkin interpolation of the resulting nodal contributions. In mathematical terms, the former involves evaluating the interaction integrand *F*(<u>x</u>), say, as: 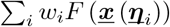 with *i* indicating the index of quadrature point, *w_i_* being the quadrature weights, and *<u>η</u>* denoting the quadrature point coordinates in the global finite element coordinate system. The latter involves the following: 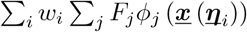 where *j* indicates the nodal index, and *φ* denotes the nodal basis functions. Based on the mathematical formulation of our method, the first integration method is more accurate. For this study we evaluated both implementations, and found minor differences in the large-scale flow structures between the two versions (*see Figure S5 for an illustration of micro-scale differences due to implementation choice*.)

### 4.4 Assumptions and limitations

The computational framework and the simulation study presented here have a few key underlying assumptions and limitations. First, we have considered a 2-dimensional geometry, and the flow-domain is in form of a channel. This assumption enabled a few advantages. The 2-dimensional computations are substantially less expensive than 3-dimensional models, thereby helping conduct a larger cohort of simulations for the systematic parametric variations we have reported here. In addition, using a simple flow-domain geometry helped control against added vortical structures originating from curvature induced secondary circulation. This ensured that any macroscale flow features and FTLE structures observed were solely due to thrombus interactions with unsteady pulsatile flow. We acknowledge that extending our simulation framework to account for 3-dimensional, and anatomically derived, geometries is a critical next step, and we are currently expanding our efforts in that direction. A second, and somewhat related, limitation is (as stated in Section 4.1) that the thrombus was treated as an already formed non-deformable aggregate. The investigation of thrombus growth was not the focus here since it is a phenomenon of considerable importance and complexity requiring a separate and dedicated discussion of its own. Within the context of a macroscale thrombus that has already formed, the no-deformation assumption needs some justification. Physiologically, thrombus aggregate formation is followed by a phase of retraction and thrombus consolidation [9,34,11], which renders a stable and compact structure to the thrombus [25]. Owing to consolidation, thus, the thrombus can be assumed to undergo small deformations at the macroscopic scale, which are negligible in comparison with other dominant flow phenomena. We also note that accounting for two-way coupled dynamic interactions between clot deformation and unsteady hemodynamics requires considerable algorithmic intricacies, and have not been addressed in this work to keep the focus on the role of flow in mediating transport processes. Furthermore, here we have assumed blood to be a Newtonian fluid, but significant variations in shear in the thrombus vicinity may lead to notable non-Newtonian effects. Extension of our hybrid particle-continuum fictitious domain finite element formulation to non-Newtonian constitutive relations is a complex task, and constitutes one of our ongoing areas of investigation. Finally, while the thrombus shapes were obtained based on experimental data [12], the microstructure and leakage values could not be equivalently derived from known experiments within the scope of this study. This was primarily due to relatively sparse availability of well characterized, fine-grained data for these parameters for integration into large-scale computations. This constitutes an area of independent investigation we are currently pursuing. Our interests include incorporating additional information on clot microstructural heterogeneities including known core-shell architectures of hemostatic plugs [44] and finer scale structural information of fibrin networks as studied in prior works [10,53]. We remark that our hybrid particle-continuum approach poses no inherent complications in terms of integrating anatomical geometries and detailed thrombus microstructure and composition data. Particularly for microstructural data, the flexibility of representing arbitrary shape and microstructure is a major advantage of our particle-continuum approach.

### 4.5 Broader implications

The computational approach and data described in this study have broader implications in terms of thrombosis disease progression and treatment. Specifically, in stroke patients thrombolytic therapy is the first course of treatment received by a patient with 3-6 hours of stroke onset [52]. Thrombolysis is often associated with poor outcomes [38] owing to incomplete understanding of drug transport in the thrombus vicinity. Precise characterization of flow-mediated transport and permeation in the thrombus environment is critical for understanding thrombolytic drug transport and subsequent thrombolytic therapy efficacy. Transport of coagulation agonists like ADP and Thromboxane released from platelets bound to the thrombus may also play a role in further thrombus deposition. Comprehensive evaluation of unsteady blood flow is also critical for assessment of flow induced loading on a thrombus and consequent thrombus fragmentation and embolization. Embolization, in particular, can cause significant complications in stroke therapy, leading to unexpected neurovascular incidents and reducing treatment efficacy. Detailed flow quantification also enables estimating local mechanical strain and shear forces. Platelets exposed to shear-loading and mechanical strain, may undergo mechanical activation, contributing further to thrombosis disease phenomena. Finally, our framework specifically addresses the need for efficient handling of arbitrary thrombus shape and microstructures representative of realistic thrombus, which is critical for advancing state-of-the-art in *in silico* analysis of thrombotic phenomena.

## 5 Concluding Remarks

We have described a parametric simulation based study on blood flow and flow-mediated transport in the neighborhood of a thrombus in arterial flow environment. The study was based on a hybrid particle-continuum fictitious domain finite element computational framework, which we have devised in prior work.

The framework enabled rapid parametric variations in thrombus microstructure for a fixed thrombus shape and size. For varying thrombus shape, microstructure, and extent of wall leakage, we evaluated: (a) the unsteady flow patterns emerging from pulsatile viscous flow interactions with the thrombus domain; (b) pressure-gradient across the thrombus boundary which drives permeation; and (c) the dynamic coherent structures identifiable from the finite time Lyapunov exponent (FTLE) field which organizes advective mass transport around the thrombus. The results from the simulation illustrated how shape, microstructure, and wall status influence flow and transport in thrombus neighborhood. The results and inferences also helped generate a unified understanding of the balance between advection, diffusion, and permeation in determining flow-mediated transport around a thrombus.

## Supporting information

Flow-Animation

FTLE-Animation

## 6 Conflicts of Interest

The Authors declare no conflicts of interest pertaining to the research presented here.

## 7 Acknowledgements

This work was partly supported by the American Heart Association (Award: 16POST27500023) and the Burroughs Wellcome Fund (Award: 1016360). This work utilized resources from the University of Colorado Boulder Research Computing Group, which is supported by the National Science Foundation (awards ACI-1532235 and ACI-1532236), the University of Colorado Boulder, and Colorado State University. The Authors also gratefully acknowledge guidance, support, and the many valuable discussions with Prof. Scott L. Diamond, Department of Chemical and Biomolecular Engineering, University of Pennsylvania. These fruitful discussions strongly benefited the study design and interpretation of results. CT performed the flow simulations, data analysis, and contributed towards manuscript content. ZI performed Lagrangian computations and data analysis. DM developed the numerical methods and computer libraries, designed the study, and wrote the manuscript. SCS contributed key inputs to finalize study design, and simulation data analysis and interpretation. All authors reviewed the manuscript and agreed to the final version.

## Notes

### Competing Interest Statement

The authors have declared no competing interest.

### Summary of Updates

Several technical details which were important for the methodology described in this manuscript have been updated; and some minor points regarding the simulation case-studies have been clarified.

