## Supplementary Materials for "Computational Investigation Of Blood Flow And Flow-mediated Transport In Arterial Thrombus Neighborhood"

### Supplementary Material

#### Animation of unsteady flow and FTLE data

Two supplementary video files have been included to visualize the dynamic evolution of the flow structures around the thrombus models with varying shape and wall leakage parameters. The first animation file, *flow-animation.mp4*, shows an animation of the unsteady flow using the colored line integration convolution (LIC) visualization as shown in Section 3.1. The second animation file, *ftle-animation.mp4*, shows an animation of the FTLE fields for the corresponding cases.

#### Supplementary data and figures

We have included here a collection of figures and quantitative data which support and supplement the test and the figures presented in the main manuscript. For each supplementary figure included here, the relevant descriptions are presented in the captions. Respective figures have been referenced at appropriate locations within the main text.

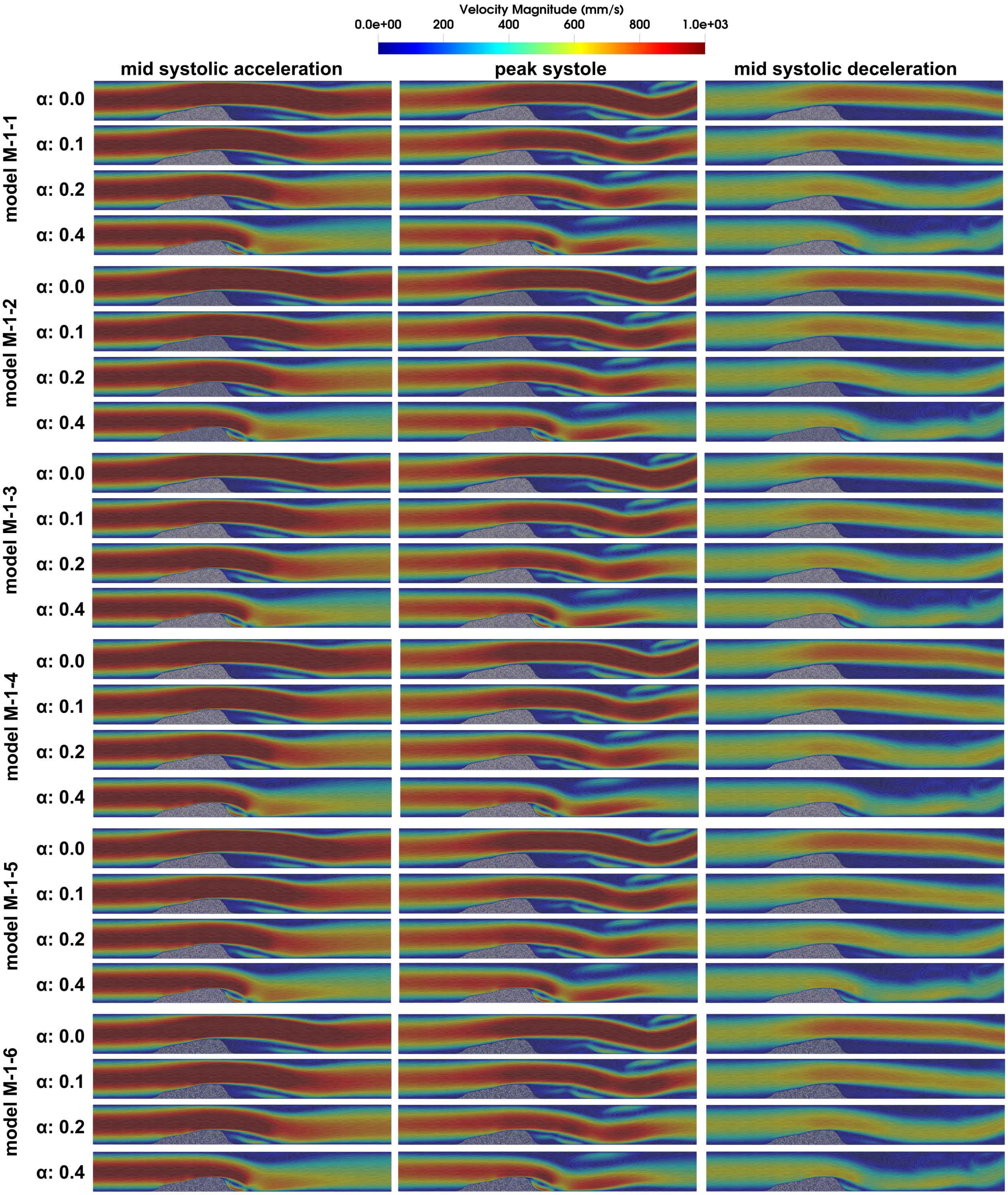


Figure S1: *Flow velocity fields at three subsequent instances in the cardiac cycle, illustrated for all combinations of thrombus model M-1-x. Figures accompany animation of flow fields presented in flow-animation.mp4.*

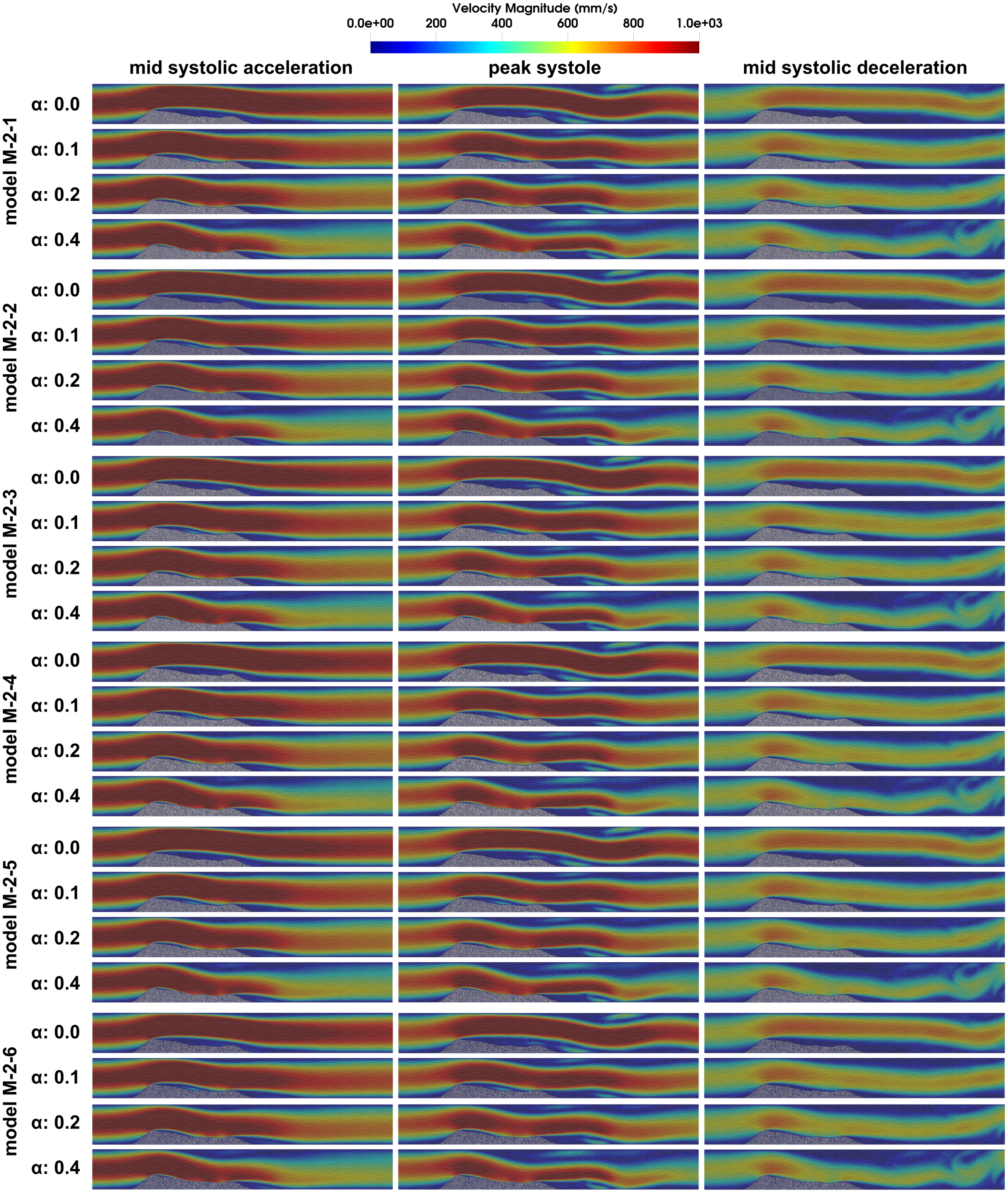


Figure S2: *Flow velocity fields at three subsequent instances in the cardiac cycle, illustrated for all combinations of thrombus model M-1-x. Figures accompany animation of flow fields presented in flow-animation.mp4.*

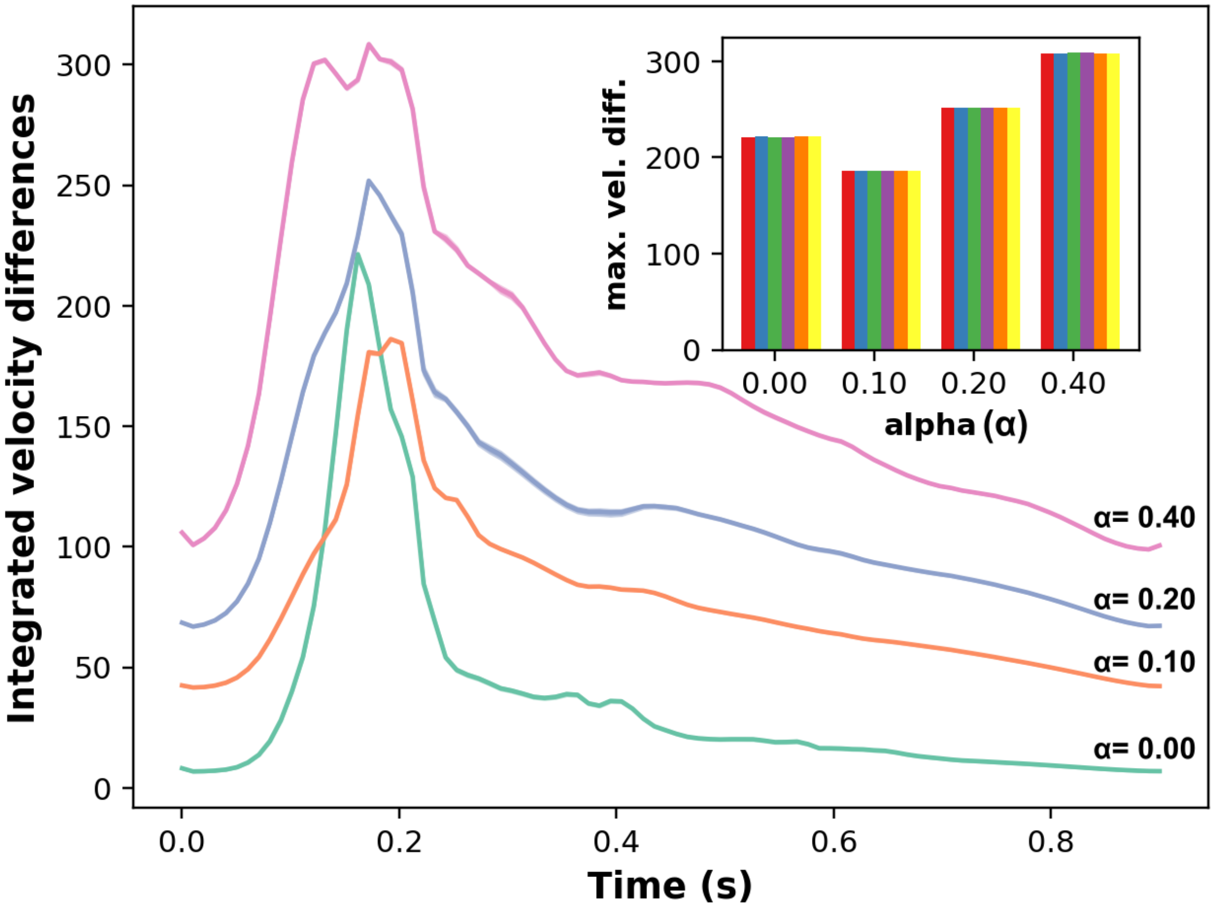


Figure S3: *Differences in flow velocity fields around models M-2-x computed with respect to flow velocities around a rigid thrombus with impermeable walls and integrated over the entire flow domain. The main visual illustrates the variation in this integrated difference over time in one cardiac cycle for varying leakage parameter $\alpha$. The curves represent mean values computed across the 6 microstructural variants. Inset visual illustrates the maximum differences for varying leakage parameters across all microstructural variants, confirming the minor influence that microstructure has in comparison with wall leakage in affecting flow around a given thrombus shape. All data are in units of velocity mm/sec.*

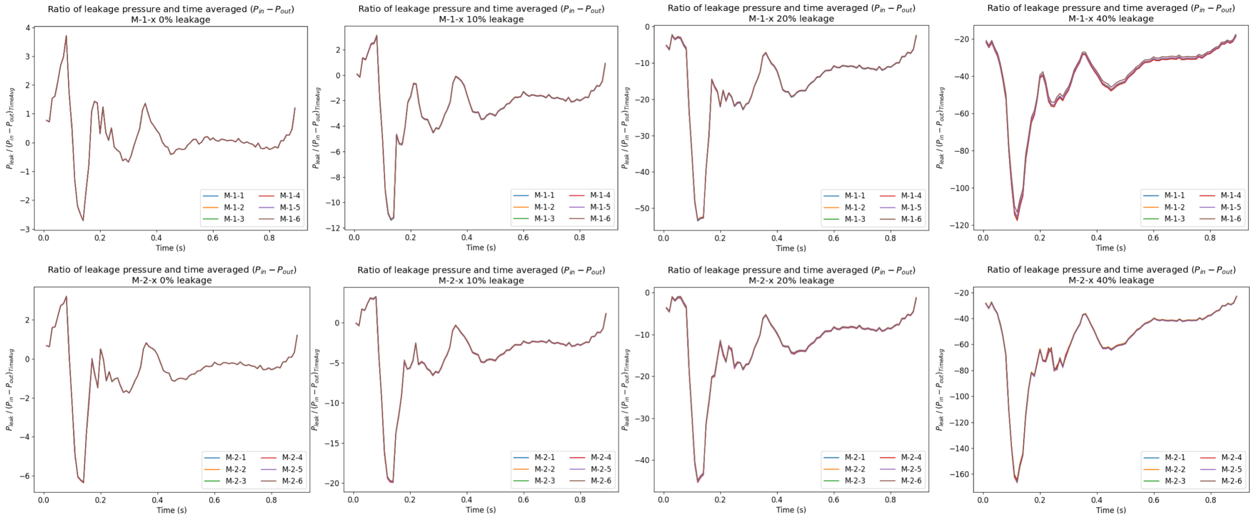


Figure S4: *Pressure at the thrombus wall leakage site, averaged across the leakage length, and scaled by the time-averaged streamwise pressure gradient. The pressure values represent the dynamic leakage flow that the simulation generates for varying extents of leakage ($\alpha$) and varying microstructures.*

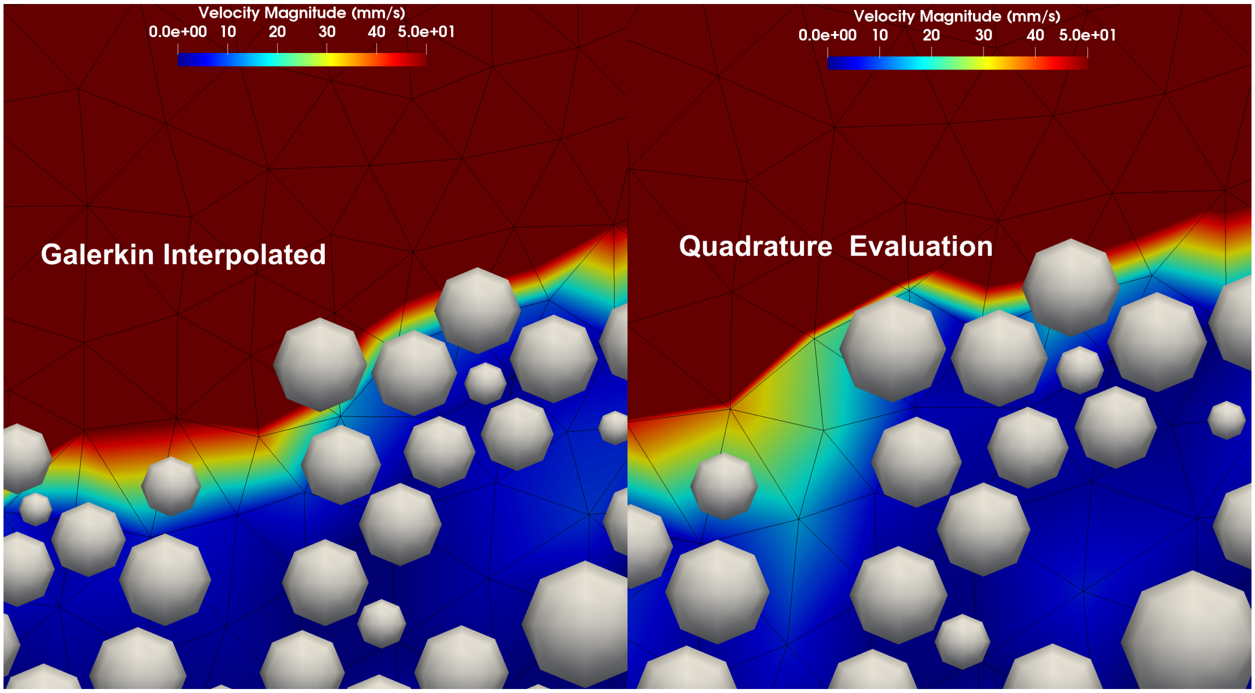


Figure S5: *A sample illustration of the small-scale differences induced due to differences between a nodal interpolation based and a quadrature point based implementation of the fictitious domain algorithm presented here. While large-scale flow structures are negligibly influenced by details of this nature, this is a factor of relevance when considering smaller scale microscopic flow features.*

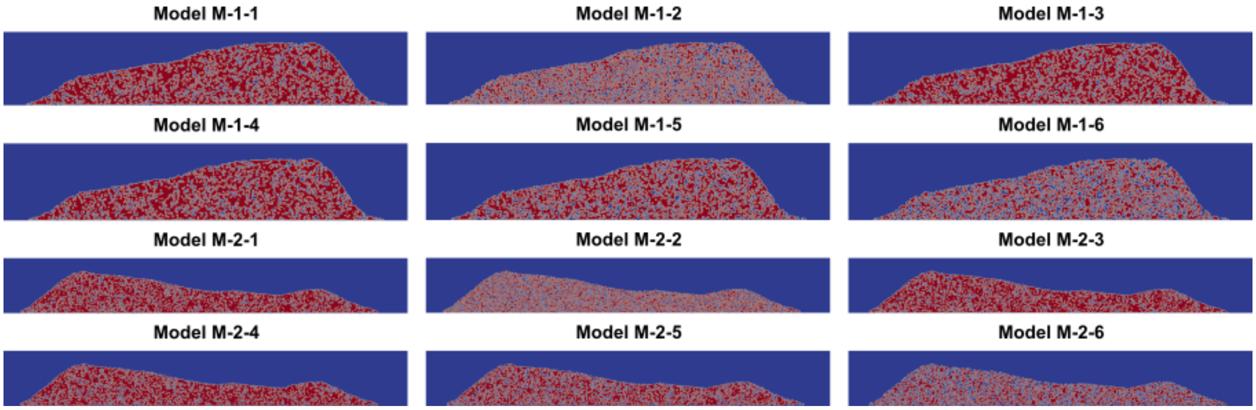


Figure S6: *Illustration of the interpolated discrete element inside-outside function onto the background fictitious domain grid. Red indicates value of 1.0 denoting 'inside' of discrete element domain, and Blue indicates 0.0 denoting 'outside' of discrete element domain. Flow occurs on the domain that is outside the discrete element domain for the thrombus.*
